# BubbleGun: Enumerating Bubbles and Superbubbles in Genome Graphs

**DOI:** 10.1101/2021.03.23.436631

**Authors:** Fawaz Dabbaghie, Jana Ebler, Tobias Marschall

## Abstract

**Motivation:** With the fast development of third generation sequencing machines, *de novo* genome assembly is becoming a routine even for larger genomes. Graph-based representations of genomes arise both as part of the assembly process, but also in the context of *pangenomes* representing a population. In both cases, polymorphic loci lead to *bubble* structures in such graphs. Detecting bubbles is hence an important task when working with genomic variants in the context of genome graphs.

**Results:** Here, we present a fast general-purpose tool, called `BubbleGun`, for detecting bubbles and superbubbles in genome graphs. Furthermore, `BubbleGun` detects and outputs runs of linearly connected bubbles and superbubbles, which we call *bubble chains*. We showcase its utility on de Bruijn graphs and compare our results to `vg`’s snarl detection. We show that `BubbleGun` is considerably faster than `vg` especially in bigger graphs, where it reports all bubbles in less than 30 minutes on a human sample de Bruijn graph of around 2 million nodes.

**Availability:** `BubbleGun` is available and documented at https://github.com/fawaz-dabbaghieh/bubble_gun under MIT license.

**Supplementary information:** Supplementary data are available at *Bioinformatics* online.

## 1 INTRODUCTION

*Genome graphs* represent collections of related sequences and have a wide range of applications in various fields of bioinformatics. In *de novo* genome assembly, for instance, graphs are used to represent a universe of plausible genome reconstructions based on a set of input sequencing reads (Miller *et al*., 2010). Recent developments have enabled even phased assembly (Porubsky *et al*., 2020), where the maternal and paternal copy of each pair of homologous chromosomes are reconstructed separately. Facilitated by GFA as an exchange data format, modern assembly tools often offer the possibility to export the underlying graphs for downstream applications. Working with graphs directly instead of using “flattened” contigs has been shown to be beneficial, for example for phased assembly (Garg *et al*., 2018), but a tool ecosystem to work with these graphs is only slowly emerging.

As a second important application domain, graphs can facilitate a comprehensive representation of genetic variation segregating in a population, called a *pangenome* (Computational Pan-Genomics Consortium, 2018). Such graph-based pangenome representations might replace present reference genomes in the future, emphasizing the need for corresponding tools.

In this work, we focus on bi-directed graphs, where sequences are represented by nodes with a left and right side. Adjacencies are then represented by edges that connect sides of two nodes and can either be non-overlapping (“blunt”) or represent an overlap between the sequences of the involved nodes. *Bubbles* are key structures within these graphs and can, for example, represent heterozygous variants in assembly graphs or polymorphisms in pangenome graphs. A subgraph between a source node *s* and a sink node *t* is defined as a superbubble (Onodera *et al*., 2013) if and only if this subgraph is directed, acyclic, and the set of nodes reachable from the source *s* is the same set of nodes from where *t* can be reached. Moreover, no other node in the superbubble should satisfy these conditions with either *s* or *t*. A *bubble* then can be defined as a special case of a superbubble, with only two disjoint paths between the source and the sink nodes (Figure 1). A linear sequence of bubbles is called a *bubble chain*. The diploid genome assembly method by Garg *et al*. (2018) highlights the importance of bubble chains. In this approach, simple bubbles reflect heterozygous variants and long reads mapped to these bubble chains in order to record which read paths in consecutive bubbles are traversed by the same reads. This gives rise to a matrix of (reads times variants) that can be used to compute a bipartition of reads into their respective haplotypes by solving the Minimum Error Correction (MEC) problem (Lippert *et al*., 2002).

**Fig. 1.**
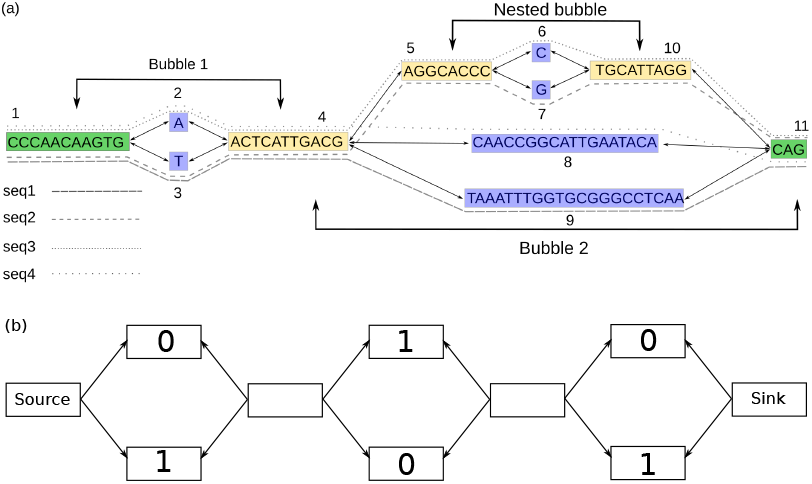
(a) A genome graphs resulting from constructing a de Bruijn graph with *k* = 9 from 4 sequences and subsequently removing overlaps (bluntification). The variances in the sequences give rise to different paths in the graph, constructing a bubble chain of one simple bubble and one superbubble with a simple bubble nested inside. (b) An example of a bubble chain with 3 bubbles, branches of bubble are randomly assigned to haplotypes 0 and 1, one haplotype takes the path going from source to sink taking only the 0 marked branches, the other haplotype takes the branches marked 1.

## 2 BUBBLEGUN

`BubbleGun` a fast general purpose tool to detect superbubbles in a given input graph by implementing the algorithm by (Onodera *et al*., 2013). In a nutshell, the algorithm iterates over all nodes *s* and determines whether there is another node *t* that satisfies the superbubble rules. `BubbleGun` can also compact linear stretches of nodes, separate biggest component of a graph, and separate user-specified neighborhood around a node for visualization and investigation. `BubbleGun` is implemented in Python and distributed as Open Source software under the terms of the MIT license.

## 3 RESULTS

### 3.1 Runtime Comparison

*Snarls* are a generalized version of superbubbles and `BubbleGun` was compared with the snarl detection algorithm (Paten *et al*., 2018) part of the `vg toolkit` (Garrison *et al*., 2018). Both tools were tested on two datasets: (1) A de Bruijn graph with a *k*-mer size of 41 representing the pangenome of 10 *Myxococcus xanthus* genomes (Supplementary Table 1) with around 600,000 nodes. (2) A de Bruijn graph with a *k*-mer size of 61 constructed from short reads from the human sample HG00733 part of the 1000 Genomes Project (1000 Genomes Project Consortium *et al*., 2015).

Time and memory consumption comparisons showed that for *M. xanthus* graph, both tools performed relatively similar with `BubbleGun` running in 20 seconds and using 0.56 Gb memory, and `VG` running in 30 seconds and using 0.85 Gb memory. However, for the HG00733 graph, `BubbleGun` took around **15 minutes** and used 22 Gb memory, where `VG` took **67 hour**s and 31 Gb memory.

### 3.2 Bubble Validation

To validate whether the bubbles detected correspond to true variants instead of repeat collapses or sequencing errors, we used a de Bruijn graph constructed from short reads data from the HG002 sample from the Genome in a Bottle (GIAB) consortium (Zook *et al*., 2016). We used a GIAB sample in order to take advantage of their high confidence variants to use for the comparison. To generate VCF files from bubble chains, we used a previously established pipeline (Ebler *et al*., 2020) that detects variants on each path separately and then merges them into a diploid VCF representation. Next, using `vcfeval` (Cleary *et al*., 2015), we compared the called variants against the high confidence variants from the HG002 sample, looking only at GIAB’s high confidence regions. This resulted in a precision of 95%. As expected, false positive bubbles are enriched in repetitive regions and when excluding regions in the repeat masker track, we observed a precision of 99%.

## 4 DISCUSSION

We presented `BubbleGun`, a tool for detecting bubbles, superbubbles, and bubble chains in genome graphs. We demonstrated that `BubbleGun` dramatically reduces the runtime of bubble detection in real world use cases, paving the way for a more widespread adoption of graph-based workflows. We expect pangenome graphs constructed from *de novo* assemblies to become a broadly used concept where traditional variant detection will be replaced by bubble detection. Diploid genome assembly (Garg *et al*., 2018) constitutes another important application area. Taken together, we envision BubbleGun to be of broad utility going forward.

## CONTRIBUTIONS

TM and FD designed the project. FD implemented BubbleGun. JE validated the bubbles. TM and FD wrote the paper.

