## Supplementary materials for "BubbleGun: Enumerating Bubbles and Superbubbles in Genome Graphs"

### BubbleGun: Enumerating Bubbles and Superbubbles in Genome Graphs. Supplementary material

| **Assembly Accession** | Bioproject | Biosample | Tax id | Species Tax id | Organism name | Infraspecific name | Version status | Assembly level | Release Type | Genome rep |
|---|---|---|---|---|---|---|---|---|---|---|
| GCA_000012685.1 | PRJNA1421 | SAMN02604018 | 246197 | 34 | Myxococcus xanthus | strain=DK 1622 | latest | Complete Genome | Major | Full |
| GCA_000278585.2 | PRJNA168264 | SAMN02471831 | 1198133 | 34 | Myxococcus xanthus | strain=DZ2 | latest | Contig | Major | Full |
| GCA_000340515.1 | PRJNA168421 | SAMN02471832 | 1198538 | 34 | Myxococcus xanthus | strain=DZF1 | latest | Contig | Major | Full |
| GCA_006400955.1 | PRJNA342411 | SAMN05757018 | 34 | 34 | Myxococcus xanthus | strain=GH3.5.6c2 | latest | Complete Genome | Major | Full |
| GCA_006401215.1 | PRJNA342411 | SAMN05757019 | 34 | 34 | Myxococcus xanthus | strain=GH5.1.9c20 | latest | Complete Genome | Major | Full |
| GCA_006401635.1 | PRJNA342411 | SAMN05757020 | 34 | 34 | Myxococcus xanthus | strain=KF3.2.8c11 | latest | Complete Genome | Major | Full |
| GCA_006402015.1 | PRJNA342411 | SAMN05757021 | 34 | 34 | Myxococcus xanthus | strain=KF4.3.9c1 | latest | Complete Genome | Major | Full |
| GCA_006402415.1 | PRJNA342411 | SAMN05757022 | 34 | 34 | Myxococcus xanthus | strain=MC3.3.5c16 | latest | Complete Genome | Major | Full |
| GCA_006402735.1 | PRJNA342411 | SAMN05757023 | 34 | 34 | Myxococcus xanthus | strain=MC3.5.9c15 | latest | Complete Genome | Major | Full |
| GCA_900106535.1 | PRJEB16561 | SAMN05444383 | 34 | 34 | Myxococcus xanthus | strain=DSM 16526 | latest | Scaffold | Major | Full |

| **Assembly Accession** | Seq Release Date | Assembly name | Submitter | Paired assembly comparison | FTP path |
|---|---|---|---|---|---|
| GCA_000012685.1 | 2006/06/07 | ASM1268v1 | TIGR | identical | ftp://ftp.ncbi.nlm.nih.gov/genomes/all/GCA/000/012/685/GCA_000012685.1_ASM1268v1 |
| GCA_000278585.2 | 2013/02/10 | ASM27858v2 | University of Iowa | identical | urlftp://ftp.ncbi.nlm.nih.gov/genomes/all/GCA/000/278/585/GCA_000278585.2_ASM27858v2 |
| GCA_000340515.1 | 2013/02/13 | Myxococcus xanthus DZF1 | University of Iowa | identical | urlftp://ftp.ncbi.nlm.nih.gov/genomes/all/GCA/000/340/515/GCA_000340515.1_Myxococcus_xanthus_DZF1 |
| GCA_006400955.1 | 2019/06/24 | ASM640095v1 | ETH Zurich | identical | urlftp://ftp.ncbi.nlm.nih.gov/genomes/all/GCA/006/400/955/GCA_006400955.1_ASM640095v1 |
| GCA_006401215.1 | 2019/06/24 | ASM640121v1 | ETH Zurich | identical | ftp://ftp.ncbi.nlm.nih.gov/genomes/all/GCA/006/401/215/GCA_006401215.1_ASM640121v1 |
| GCA_006401635.1 | 2019/06/24 | ASM640163v1 | ETH Zurich | identical | ftp://ftp.ncbi.nlm.nih.gov/genomes/all/GCA/006/401/635/GCA_006401635.1_ASM640163v1 |
| GCA_006402015.1 | 2019/06/24 | ASM640201v1 | ETH Zurich | identical | urlftp://ftp.ncbi.nlm.nih.gov/genomes/all/GCA/006/402/015/GCA_006402015.1_ASM640201v1 |
| GCA_006402415.1 | 2019/06/24 | ASM640241v1 | ETH Zurich | identical | urlftp://ftp.ncbi.nlm.nih.gov/genomes/all/GCA/006/402/415/GCA_006402415.1_ASM640241v1 |
| GCA_006402735.1 | 2019/06/24 | ASM640273v1 | ETH Zurich | identical | urlftp://ftp.ncbi.nlm.nih.gov/genomes/all/GCA/006/402/735/GCA_006402735.1_ASM640273v1 |
| GCA_900106535.1 | 2016/10/22 | IMG-taxon 2693429903 | DOE | identical | urlftp://ftp.ncbi.nlm.nih.gov/genomes/all/GCA/900/106/535/GCA_900106535.1_IMG-taxon_2693429903_annotated_assembly |

Supplementary Table. 1: Information on the 10 Myxococcus *xanthus* used for testing and results

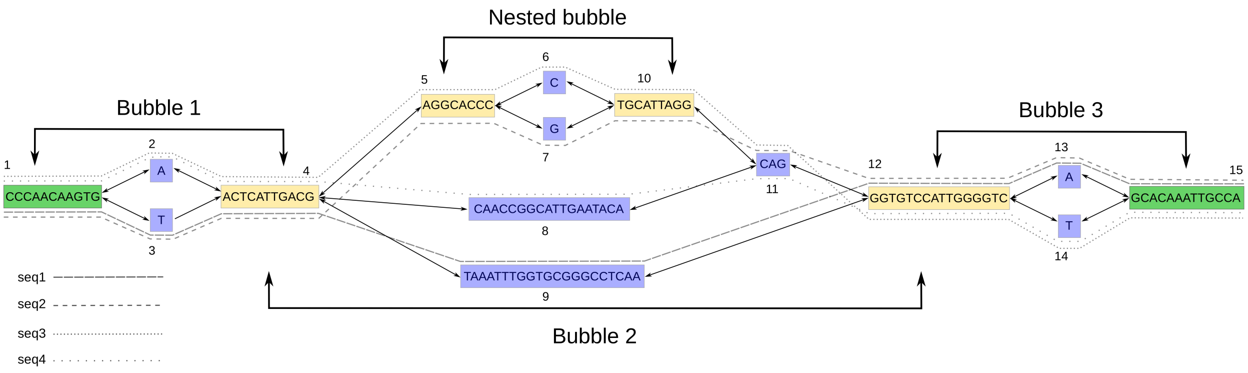


Supplementary Figure. 1: Bluntified de Bruijn graph constructed from 4 sequences presented in Supplementary Table

### 1 More on methods

#### 1.1 Bulding de Bruijn Graphs

First, short reads were corrected using `Lighter` (Song *et al.*, 2014). Afterwards, `bcalm2` (Chikhi *et al.*, 2016) was used to build the de Bruijn graphs from the corrected reads. For the two data sets used for testing, *M. xanthus* and the human HG00733 short reads, a $k$-mer minimum abundance cutoff was set to 2, the rest of the arguments were left to default, and for the bubbles validation step, the $k$-mers minimum abundance cutoff was set to 3, as the reads here had higher coverage.

#### 1.2 Algorithm

The algorithm presented by Onodera *et al.* (2013) keeps a dynamic set $S$ that contains the nodes that are possible to visit in the next iterations, once one of these node is visited, it is popped from $S$. It also keeps track of visited and seen nodes, where seen nodes are nodes that have at least one visited parent. The algorithm aborts when finding a node with no children (a tip) or an edge that points back to $s$ (a cycle). Once there is only one node $t$ left in $S$ and no other nodes are marked as seen the algorithm then returns $t$ as the sink of the superbubble.
The algorithm here expects a bi-directed graph. Therefore, for every node we start from, we can go in two directions, e.g. in Supplementary Figure 1, if we start at node 4, we can start looking to both ‘left’ and ‘right’ of the graph, however, when entering a node from one side we have to exit from the opposite side to keep the directionality. To chain bubbles and superbubbles together, we can simply call the same algorithm recursively on the returned node $t$, and continue the search from the opposite direction where we entered $t$, e.g. in Figure 1 in paper, starting from node 4 going ‘right’ the algorithm will return

| | |
|---|---|
| Sequence_1 | CCCAACAAGTGTACTCATTGACGTAAATTTGGTGCGGGCCTCAAGGTGTCCATTGGGGTCAGCACAAATTGCCA |
| Sequence_2 | CCCAACAAGTGTACTCATTGACGAGGCACCCGTGCATTAGGCAGGGTGTCCATTGGGGTCTGCACAAATTGCCA |
| Sequence_3 | CCCAACAAGTGAACTCATTGACGAGGCACCCCTGCATTAGGCAGGGTGTCCATTGGGGTCAGCACAAATTGCCA |
| Sequence_4 | CCCAACAAGTGAACTCATTGACGCAACCGGCATTGAATACACAGGGTGTCCATTGGGGTCTGCACAAATTGCCA |

Supplementary Table. 2: Sequences used to generate the example in Supplementary Figure 1 and Figure 1

node 12 as the sink of the superbubble, calling the algorithm again on node 12 and looking to the 'right' will return node 15. Once the algorithm aborts without returning a node, we go back to node 4 and keep looking to the 'left' until the algorithm aborts again. The result of this will be a chain of bubbles and superbubbles. Moreover, as the graph is compacted (no stretch of several nodes with in and out degree of one exist), we know that a chain of bubbles would share sources and sinks, i.e. the source of one bubble can be the sink of another. Therefore, this guarantee that calling the algorithm recursively would find the next bubble if it exists.

##### 1.3 Nested bubbles

To detect nested bubbles inside of superbubbles and report that relation, the algorithm was called recursively on on each superbubble and any chain (whether it consists of only simple bubbles, superbubbles, or both) will be reported as having a parent superbubble and parent chain, and the respective IDs will be reported out in the JSON output.

#### 2 Bubbles Validation

To find out if a bubble represent a variant in the genome, and where in the genome this variant is located, we extracted two sequences from every bubble chain, where a sequence is the concatenation of the node labels - with overlaps removed - of a path going from one end of the chain to the other. The two sequence represent two randomly-chosen haplotypes, Figure 1 (b) in paper.
We then aligned these haplotypes back to the reference genome using `minimap2` (Li, 2018), called variants separately on each haplotype using the `paftools.js` script with the `minimap2` package and then merged the variants into a diploid VCF representation, using a similar pipeline to call variants presented in the tool `PanGenie` (Ebler *et al.*, 2020). The command used for separating the two haplotypes using `BubbleGun` is : `BubbleGun -g hg002_short_reads.gfa -k 61 bchains --only_simple --out_haplos`. This will output two FASTA files named `haplotype1.fasta` and `haplotype2.fasta`.
The pipeline used to align these two FASTA files back to the reference and calling the variance can be found here `https://bitbucket.org/jana_ebler/vcf-merging/src/chains-genotyping/`

#### 2.1 False Positives Assessment

After comparing the variants called from the bubble chains with the high confidence variants provided by the 1000 Genomes Consortium for the HG002 samples, we found that 5% were false positive, after filtering against repeat regions, we lower the number to around 1%. Looking more in detail to the left 1% false positive variants we see mostly two causes:

1. Positions in the genome where 3 or more raw reads had the same error. As `bcalm2` was set to an abundance minimum of 3, which means that if the $k$-mer appears in 3 reads or more, it will be included in the graph. Figure 2

2. Alignment problems towards the end of a haplotype alignment. This can also be seen when aligning long reads sometimes. Figure 3 is an example.

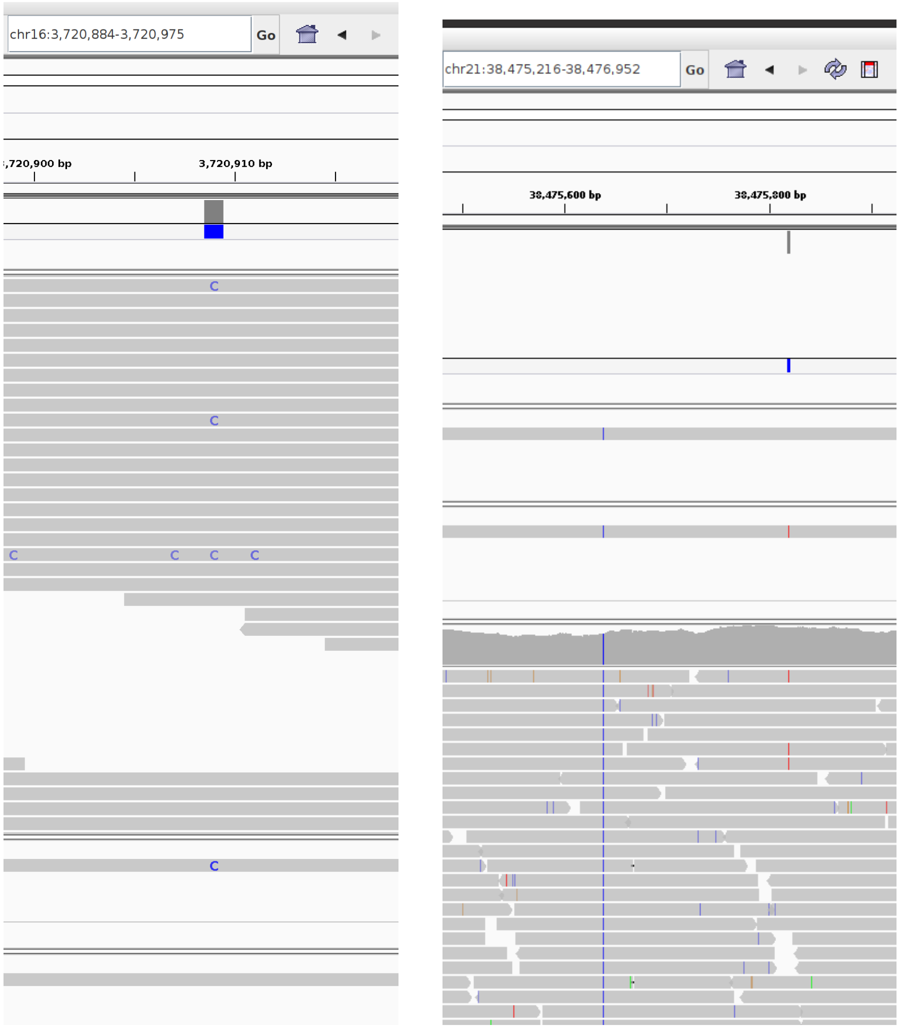


Supplementary Figure. 2: Two screenshots from IGV (Interactive Genome Viewer) showing the false positive variant and that 3 raw reads containing that variant at that position

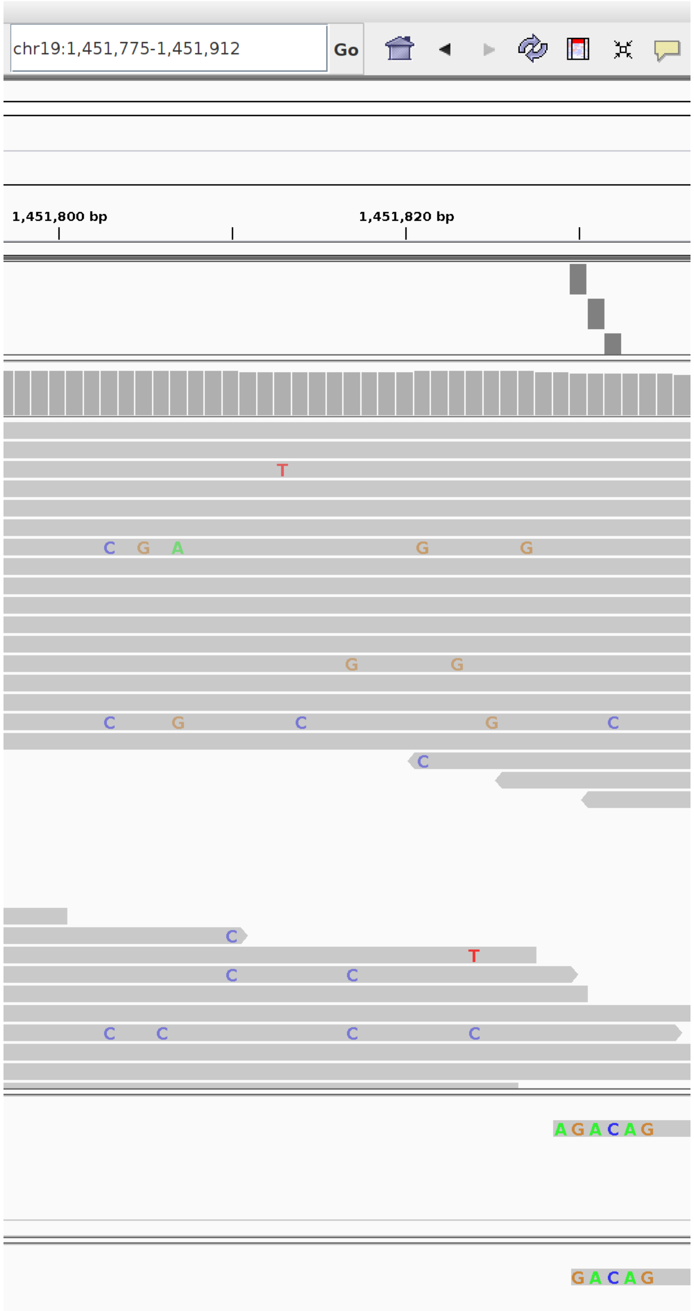


Supplementary Figure. 3: The two bottom tracks are the two haplotypes aligned, and we see that the first 5-6 amino acids of the haplotypes are erroneous, and could be caused because of the highly erroneous short reads in this section of the alignment
